# Temperature-regulated *FLOWERING LOCUS T* homologs make distinct contributions to the floral transition in vernalization-dependent and vernalization-independent *Taraxacum koksaghyz* plants

**DOI:** 10.1101/2025.03.03.640718

**Authors:** Andrea Känel, Kai-Uwe Roelfs, Michael Wissing, Benjamin Lenzen, Malin Klein, Richard M. Twyman, Gundula A. Noll, Dirk Prüfer

**Author notes:** Corresponding author: Andrea Känel.

## Abstract

Flowering that requires a period of cold (vernalization) is a key trait in many crops, but the underlying regulatory pathways are often poorly understood. *Taraxacum koksaghyz* is a rubber-producing dandelion of the family Asteraceae, which also includes other economically important crops such as chicory and lettuce. Most *T. koksaghyz* plants require cold exposure to induce flowering, but plants that have lost the dependence on vernalization are more suitable for domestication and breeding. To provide insight into the molecular basis of mandatory vernalization in *T. koskaghyz*, we identified three *FLOWERING LOCUS T* (*FT*) homologs (*TkFT1–3*) that are differentially expressed under varying environmental conditions. *TkFT1* and *TkFT2* are expressed under long-day conditions at ambient temperatures whereas only *TkFT1* is weakly expressed under short-day conditions. Exposure to cold was shown to repress *TkFT1* but induce *TkFT3*. Overexpression experiments revealed that *TkFT1–3* bypass the vernalization requirement in *T. koksaghyz* and its close relative *T. officinale*, and promote early flowering in vernalization-independent *T. brevicorniculatum*. We also identified two *FRUITFULL* homologs (*TkFUL1* and *TkFUL2*) as downstream targets, which were upregulated in *TkFT* overexpression lines. Our findings suggest that TkFT1 promotes vernalization-independent flowering, whereas *TkFT3* expression during the cold period is needed to promote vernalization-dependent flowering. This study explicates the regulatory network controlling flowering time in *T. koksaghyz*, contributing to a broader understanding of flowering in the family Asteraceae and providing knowledge that can be used in the future to facilitate domestication and breeding.

**Key message:** In *Taraxacum koksaghyz*, FT homologs function as floral inducers that upregulate *FRUITFULL* homologs, with TkFT1 linked to vernalization-independent flowering and TkFT3 associated with the acquisition of flowering competence during vernalization.

## Introduction

The Russian dandelion (*Taraxacum koksaghyz*) is a diploid, perennial plant species belonging to the family Asteraceae. It is a promising alternative source of high-quality natural rubber, as well as other secondary metabolites such as inulin and triterpenes (van Beilen and Pourir 2007; Schulze Gronover et al. 2011; Pütter et al. 2019). It can be cultivated in temperate climates and thrives on nutrient-poor soils that cannot be used to cultivate other crops, therefore providing a sustainable alternative to the Pará rubber tree (*Hevea brasiliensis*), which is restricted to tropical regions and requires specific soil types (Javorvsky 1944; van Beilen and Pourir 2007). However, the domestication of *T. koksaghyz* is hindered by traits such as self-incompatibility (which leads to high heterozygosity) and the need for most plants to undergo vernalization (exposure to prolonged cold temperatures) to achieve synchronized and vigorous flowering, a prerequisite for efficient seed harvesting over a short period of time (Whaley and Bowen 1947; Suomela 1950; Krotkov 1945). A deeper understanding of the exogenous and endogenous factors that regulate flower development is therefore needed to facilitate the breeding of vernalization-independent (VI) cultivars with shorter life cycles and improved seed production, making *T. koksaghyz* economically viable for the sustainable production of natural rubber.

Flowering is intricately regulated by both endogenous and environmental factors, including hormonal signaling, plant age, temperature, and photoperiod, ensuring that floral development occurs under favorable conditions. The underlying mechanisms have been extensively studied in Arabidopsis (*Arabidopsis thaliana*), the primary model organism for flower development. Briefly, in winter-annual Arabidopsis, the transcription factor FLOWERING LOCUS C (FLC) inhibits flowering by repressing the expression of key floral integrator genes, such as *FLOWERING LOCUS T* (*FT*), *SUPPRESSOR OF OVEREXPRESSION OF CONSTANS 1* (*SOC1*), and *FD*, the latter encoding a bZIP transcription factor that acts as a cofactor of FT (Hepworth et al. 2002; Abe et al. 2005; Helliwell et al. 2006; Searle et al. 2006). Vernalization leads to the suppression of *FLC* expression by ensuring persistent histone methylation at the *FLC* locus (Sheldon et al. 2000; Bastow et al. 2004). Day length also plays a pivotal role in the regulation of flowering in Arabidopsis because CONSTANS (CO), which activates the expression of *FT*, is only stabilized under long-day (LD) conditions (Fornara et al. 2009). The FT protein, which belongs to the PHOSPHATIDYLETHANOLAMINE-BINDING PROTEIN (PEBP) family, is mainly produced in the leaves and is transported via the phloem to the shoot apical meristem (SAM), where it forms a florigen activation complex by interacting with FD, mediated by 14-3-3 proteins (Wigge et al. 2005; Corbesier et al. 2007; Taoka et al., 2011). The FT paralog TERMINAL FLOWER 1 (TFL1) also interacts with FD and 14-3-3 proteins, together forming the florigen repressor complex. FT and TFL1 are assumed to compete for interaction with FD, resulting in either activation or repression of their shared target genes, such as *FRUITFULL* (*FUL*), *LEAFY* (*LFY*) and its downstream target *APETALA1* (*AP1*) (Zhu et al. 2020). Activation eventually leads to flowering (Weigel and Nilson, 1995; Liljegren et al. 1999; Ferrándiz et al. 2000).

The genetic pathways regulating floral transition and flowering time have been investigated in species other than Arabidopsis, revealing substantial divergence and adaptation (Blümel et al. 2015). Although the perception of signals such as vernalization and day length can be conveyed by proteins other than FLC or CO, as is the case in sugar beet (*Beta vulgaris*) (Pin et al. 2012; Dally et al. 2018) and temperate cereals (Ream et al. 2012), the FT protein appears to be an important network hub that is conserved between species (Jin et al. 2021). Arabidopsis FT is antagonized by TFL1, but in several other species, such as sugar beet, tobacco (*Nicotiana tabacum*), and soybean (*Glycine max*), antagonistic FTs have evolved as a means to balance the regulation of flowering time (Pin et al. 2010; Harig et al. 2012; Liu et al. 2018).

Although some modern crops such as chicory and endive (*Cichorium* spp.), garden lettuce (*Lactuca sativa*), and sunflower (*Helianthus annuus*) are members of the family Asteraceae, little is known about the molecular control of floral transition in this family (Leijten et al. 2018). However, *FT* genes appear to play an important role in several Asteraceae species. For example, differences in the sequence and expression of sunflower FT proteins and their interaction with co-factors have played a key role in sunflower domestication (Blackman et al. 2010). In the ornamental crop *Chrysanthemum*, the interplay between FT and TFL1 proteins confers the obligate short-day (SD) flowering phenotype (Oda et al. 2012; Higuchi et al. 2013). In lettuce, the floral activator LsFT may promote unwanted heat-induced bolting (Fukuda et al. 2011; Chen et al. 2017).

In the facultative LD plant *T. koksaghyz*, most but not all individuals show vernalization-dependent (VD) flowering (Krotkov 1945, Borthwick et al. 1943). Three major loci with epistatic interactions may govern the dependence on vernalization (Hodgson-Kratky and Wolyn 2015), highlighting the complexity of the floral regulatory network in this species. Recently, we used whole-genome bisulfite sequencing (WGBS) in *T. koksaghyz* to investigate epigenetic variation in VI and VD plants, and combined this with gene expression data acquired using the massive analysis of cDNA ends (MACE) technique. We compared VI and VD plants before, during and after vernalization, revealing several promising floral candidate genes (Roelfs et al. 2024), but the role of *FT* genes remains to be elucidated.

Here, we characterized three *T. koksaghyz FT* paralogs (*TkFT1–3*) to determine their expression profiles under different environmental conditions. We also conducted functional analysis by overexpression and interaction analysis with TkFD1 to find out whether TkFT1–3 are components of the *T. koksaghyz* florigen activation complex. We identified *FUL* genes in several *Taraxacum* species and provide evidence that they are downstream targets of TkFT1–3. Our findings offer insights into the complex regulatory network governing flowering in *T. koksaghyz* and will be useful for future domestication and breeding programs for this alternative rubber crop.

## Materials and Methods

### Plant material and cultivation conditions

We obtained *T. brevicorniculatum* and *T. officinale* seeds from the Botanical Garden Karlsruhe (Germany) and the Botanical Garden Münster (Germany), respectively. *T. koksaghyz* plants were provided by ESKUSA GmbH (Straubing, Germany). *T. koksaghyz* plants Tk203, TkVD.1, TkVD.3, TkVI.1 and TkVI.3 were selected from a wild-type mixed population and phenotyped for 200 days in the greenhouse. TkVI.1 and TkVI.3 flowered without vernalization within 58 days, whereas Tk203, TkVD.1 and TkVD.3 did not flower without vernalization within 200 days. Tk203, TkVD.1, TkVD.3, TkVI.1 and TkVI.3 plants were propagated *in vitro* to obtain clones as previously described but without antibiotic selection (Stolze et al. 2017). If not stated otherwise, plants were cultivated in a climate-controlled greenhouse at 14–18 °C (night) and 22–25 °C (day) with a 16-h photoperiod (artificial light switched on if natural light fell below 700 μmol m^−2^ s^−1^). For diurnal expression analysis, plants were cultivated under LD conditions in the greenhouse as described above or under SD conditions (8-h photoperiod, 200 μmol m^−2^ s^−1^, 22 °C under light, 20 °C in the dark). Plants were vernalized in a Percival LT-36VL phytochamber (CLF Plant Climatics, Wertingen, Germany) at 6 °C (8-h photoperiod, 200 μmol m^−2^ s^−1^) for the indicated time. Plant material and cultivation conditions for expression analysis by MACE were described in detail in our previous study (Roelfs et al. 2024).

### Sequence verification and cloning

RNA was isolated from *T. koksaghyz* leaves or SAM-enriched tissue using the innuPREP Plant RNA kit (IST Innuscreen, Berlin, Germany) and reverse-transcribed using the PrimeScript RT master mix (Takara Bio, San Jose, CA, USA). *TkFT1–3*, *TkFD1*, *TkFUL1* and *TkFUL2* coding sequences were amplified from cDNA using Phusion high-fidelity polymerase (Thermo Fisher Scientific, Waltham, MA, USA) and the primers listed in Table S1, transferred to pCRII-Topo (Thermo Fisher Scientific) and sequenced. Lettuce genomic DNA isolated using the NucleoSpin Plant II Kit (Machery Nagel, Düren, Germany) was used to amplify the polyubiquitin promoter (PLsUbi) using the primers listed in Table S1, and splicing by overlap extension (SOE) PCR was used to disrupt the internal XbaI site. The PCR product with attached restriction sites was transferred to vector pLab12.1 (Post et al. 2012). *TkFT1–3* overexpression constructs were prepared by reverse-transcribing RNA from *T. koksaghyz* Tk203 leaves using PrimeScript RT master mix. The *TkFT1–3* cDNAs were amplified with attached restriction sites (Table S1) and transferred to pLab12.1PLsUbi by restriction and ligation. For BiFC assays, *TkFT1–3* and *TkFD1* cDNAs were amplified with attached restriction sites (Table S1) and transferred to pENTR4 (Thermo Fisher Scientific) by restriction and ligation. Subsequent transfer to pBatTL vectors was achieved by Gateway recombination using Gateway LR clonase II (Thermo Fisher Scientific). The pBatTL plasmids were kindly provided by Joachim Uhrig and Guido Jach (University of Cologne, Cologne, Germany).

### Quantitative real-time PCR

Plant tissues were harvested, snap-frozen in liquid nitrogen and ground using either an MM400 bead mill (Retsch, Haan, Germany) or with a mortar and pestle under liquid nitrogen. RNA was extracted using the innuPREP Plant RNA kit. Residual genomic DNA was digested using the TURBO DNA-free kit (Thermo Fisher Scientific). RNA was reverse-transcribed using PrimeScript RT master mix and gene expression was analyzed by quantitative real-time PCR (qRT-PCR) using the Kapa SYBR Fast qPCR Master Mix (Merck, Darmstadt, Germany), the primers listed in Table S1, and the CFX96 Real-Time System (Bio-Rad Laboratories, Hercules, CA, USA). Each reaction was carried out in technical triplicates. Specificity was ensured by melt curve analysis, by including no-template and no-reverse-transcription controls, and the sequencing of PCR products. Individual PCR efficiency was determined using LinReg PCR v2017.0 and relative gene expression levels were normalized to the *RP* gene.

### *Agrobacterium*-mediated transformation

All *Taraxacum* species were transformed using *Agrobacterium tumefaciens* as previously described (Stolze et al. 2017). The transgenic plants were selected by cultivation on medium containing phosphinothricin. Transgene integration was confirmed by isolating genomic DNA followed by PCR using the KAPA 2G robust PCR kit (Merck) and the primers listed in Table S1.

### Bimolecular fluorescence complementation assays

Bimolecular fluorescence complementation (BiFC) assays were carried out following the infiltration of *Nicotiana benthamiana* leaves and the analysis of leaf disks as previously described (Mäckelmann et al. 2024).

### Identification of PEBP-like proteins

The hidden Markov model (HMM) of the PEBP family (PF01161) was downloaded from the Pfam database (Paysan-Lafosse et al. 2025) and used to screen the predicted protein sequences of the reference genome (Lin et al. 2022) using the *hmmsearch* command in HMMER v3.4 (Eddy 2011) with an E-value cutoff of < 0.001.

### Phylogenetic analysis

PEBP-like proteins other than those identified in this study were adapted from Harig et al. (2012). Multiple sequences were aligned using MEGA v11.0.13 (Tamura et al. 2021) with the MUSCLE algorithm. The approximately-maximum-likelihood tree was constructed using FastTree v2.1 (Price et al. 2010) with the gamma option. The tree was visualized using iTOL v7.0 (Letunic and Bork 2024).

### MACE data analysis

MACE sample preparation, processing, quality control and sequencing were carried out as previously described (Roelfs et al. 2024). Reads were mapped to the *T. koksaghyz* reference genome (Lin et al. 2022) and quantified using subread-align (maximum of five best-quality alignments reported) and featureCounts (multi-mapping reads counted fractionally) in Subread v2.0.3 (Liao et al. 2013). Raw counts were analyzed for differential expression using edgeR v3.40.2 (Chen et al. 2016). We calculated p-values and false discovery rates (FDRs) using a minimum of log_2_(1.5). For each comparison, genes were considered differentially expressed if the FDR was < 0.05.

### Analysis of promoters *in silico*

Approximately 1.6 kb of the *TkFT1* and *TkFT3* promoter sequences was amplified using the primers listed in Table S1 and genomic DNA isolated from the VD accession Tk203. The PCR products were transferred to pCRII-Topo and sequenced. The promoter sequences were uploaded to the promoter analysis tool of the PlantPan 4.0 sub-database PCBase 2.0 (Chow et al. 2024) with Arabidopsis as the reference organism. Transcription factor binding site (TFBS) hits were filtered to exclude those with low similarity (< 0.8). PCBase 2.0 integrates experimental data (e.g. ChIP-Seq) to find transcription factors that bind TFBSs in Arabidopsis. Gene Ontology (GO) terms for the identified transcription factors were categorized using the GO enrichment tool of The Arabidopsis Information Resource (TAIR) (www.arabidopsis.org). The data were visualized using ggplot2 v3.5.1 (Wickham 2016) and patchwork v1.3.0 (Pedersen 2024).

### Statistics

If not stated otherwise, all bar plots were generated in OriginPro 2023 (OriginLab Corporation, Northampton, MA, USA). With the exception of MACE data, statistical analysis was carried out in OriginPro 2023. Equality of variances was determined by one-way analysis of variance (ANOVA), and pairwise comparisons were assessed using Tukey’s *post hoc* test. Spearman rank correlation was determined using two-sided tests.

## Results

### Identification of *TkFT1–3*

Seven putative *FT* genes were identified in the published *T. koksaghyz* genome (Lin et al. 2022) by using the PEBP family HMM profile to search predicted protein sequences. Plant PEBP family proteins can be assigned to three main clades: FT-like, MFT-like and TFL-like (Chardon and Damerval, 2005). *T. koksaghyz* PEBPs were integrated into this classification by building an approximately-maximum likelihood tree from a multiple sequence alignment of the *T. koksaghyz* PEBP sequences and others previously characterized by Amaya et al. (1999), Karlgren et al. (2011), and Harig et al. (2012). Two of the *T. koksaghyz* proteins fitted into the MFT-like clade, three into the TFL-like clade and two into the FT-like clade (Fig. 1). Based on their phylogenetic relationship to Arabidopsis homologs, the MFT-like proteins were named TkMFT1 and TkMFT2, the TFL-like proteins were named TkBFT1, TkBFT2 and TkTFL1, and the two FT-like proteins were named TkFT1 and TkFT2.

**Fig. 1.**
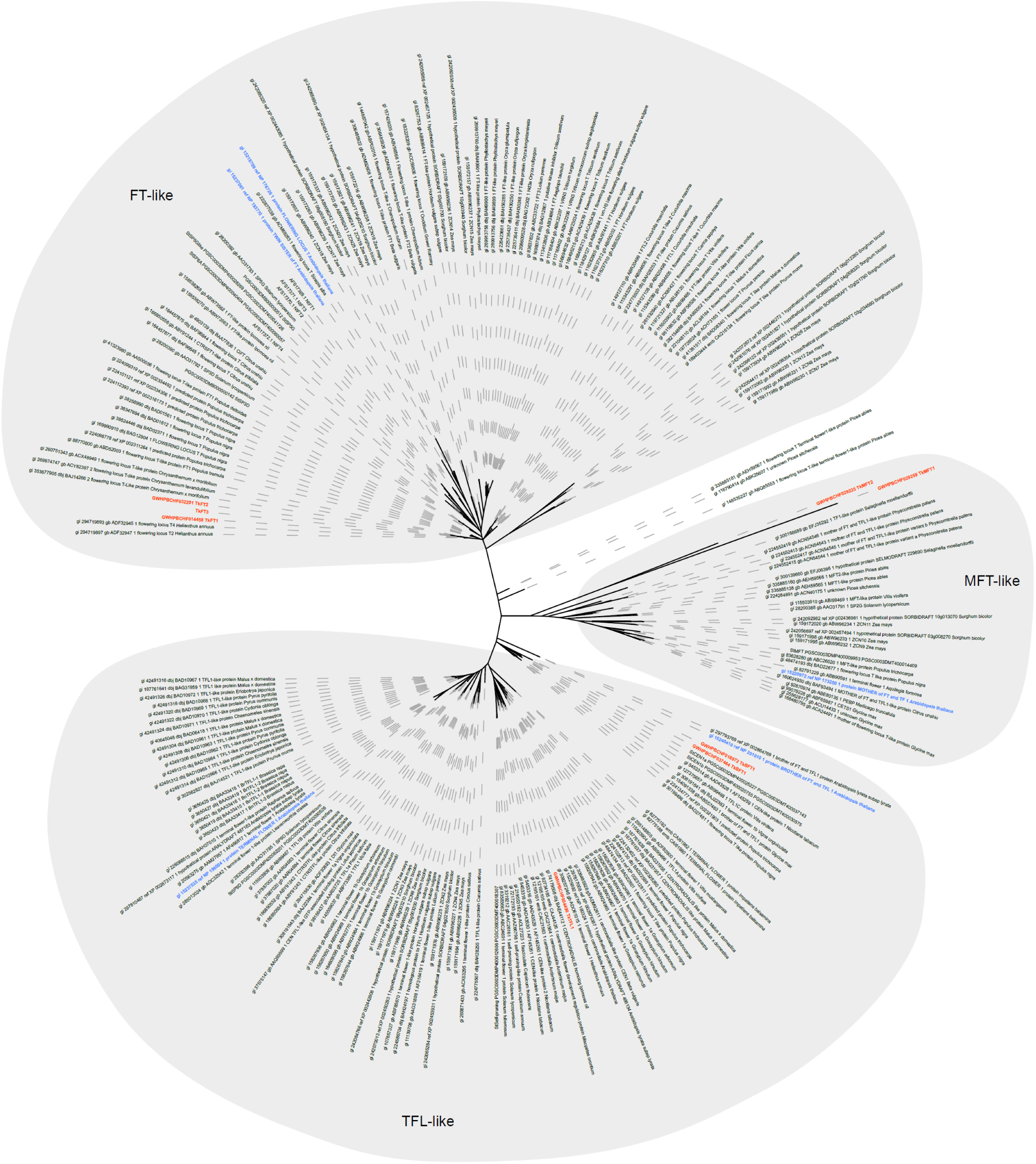
Phylogenetic tree of the plant PEBP family defined by Amaya et al. (1999), Karlgren et al. (2011) and Harig et al. (2012), including the PEBPs from *T. koksaghyz*. Representative Arabidopsis homologs are shown in blue and *T. koksaghyz* PEBPs are shown in red. FT = FLOWERING LOCUS T, TSF = TWIN SISTER OF FT, MFT = MOTHER OF FT AND TFL 1, TFL = TERMINAL FLOWER, BFT = BROTHER OF FT AND TFL 1.

Next, we amplified the *TkFT1* and *TkFT2* coding sequences by reverse-transcribing RNA isolated from the leaves of VI and VD *T. koksaghyz* plants. Sequence analysis of the cDNAs revealed full-length open reading frames (ORFs) for both genes. Surprisingly, primers for the amplification of *TkFT2* generated an additional ORF whose deduced amino acid sequence fitted into the FT-like clade. We named this ORF *TkFT3* (Fig. 1). At the amino acid level, TkFT1–3 showed 91.3–93.1% sequence identity to each other and 70.7–74.3% sequence identity to Arabidopsis FT.

### *TkFT1* and *TkFT3* are differentially expressed during vernalization

VD *T. koksaghyz* plants require approximately 2 weeks of cold exposure to reliably induce flowering within a short time frame (Fig. S1A). To gain more insight into the expression levels of *TkFT1–3* in VI and VD plants under the conditions needed for floral induction, we reanalyzed our MACE data from leaves of VI and VD plants before, during and after vernalization, and from SAM-enriched tissue during reproductive and vegetative growth (Roelfs et al. 2024). We found that *TkFT1* and *TkFT3* were differentially expressed in the leaves of VD plants during vernalization (Fig. 2A, C). Although *TkFT1* was expressed in VD plants before and after vernalization, its expression level decreased sharply during vernalization. In contrast, *TkFT3* expression was only induced during vernalization. *TkFT1* was strongly expressed in VI plants and, to a lesser extent, also in VD plants before vernalization, whereas *TkFT3* is not expressed in VI plants and only minimally in VD plants before vernalization. Both genes were expressed at low levels in SAM-enriched tissue (Fig. 2A, C). *TkFT2* was expressed at rather low levels in VI and VD plants before, during and after vernalization, but was not expressed in the SAM-enriched tissue (Fig. 2B). Remarkably, none of the genes showed a significant increase in expression after vernalization (Fig. 2A-C). The expression levels determined by MACE were verified by qRT-PCR using pooled material from corresponding leaf samples, confirming that *TkFT1* is downregulated and *TkFT3* is upregulated during vernalization. We also verified general gene expression level tendencies (Fig. S2). To determine whether *TkFT1–3* expression profiles differ between VI and VD plants during vernalization, we measured their expression levels by qRT-PCR in VI and VD plants before, during and after vernalization. The high level of *TkFT1* expression in VI plants, and the lower level in VD plants, were downregulated to minimal levels during vernalization (Fig. 2D). *TkFT2* expression was inconsistent in the plants we analyzed, increasing in one of the VD accessions during vernalization but remaining rather low in the others (Fig. 2E). *TkFT3* expression was induced in both VI and VD plants, but expression levels were higher in VD plants (Fig. 2F).

**Fig. 2.**
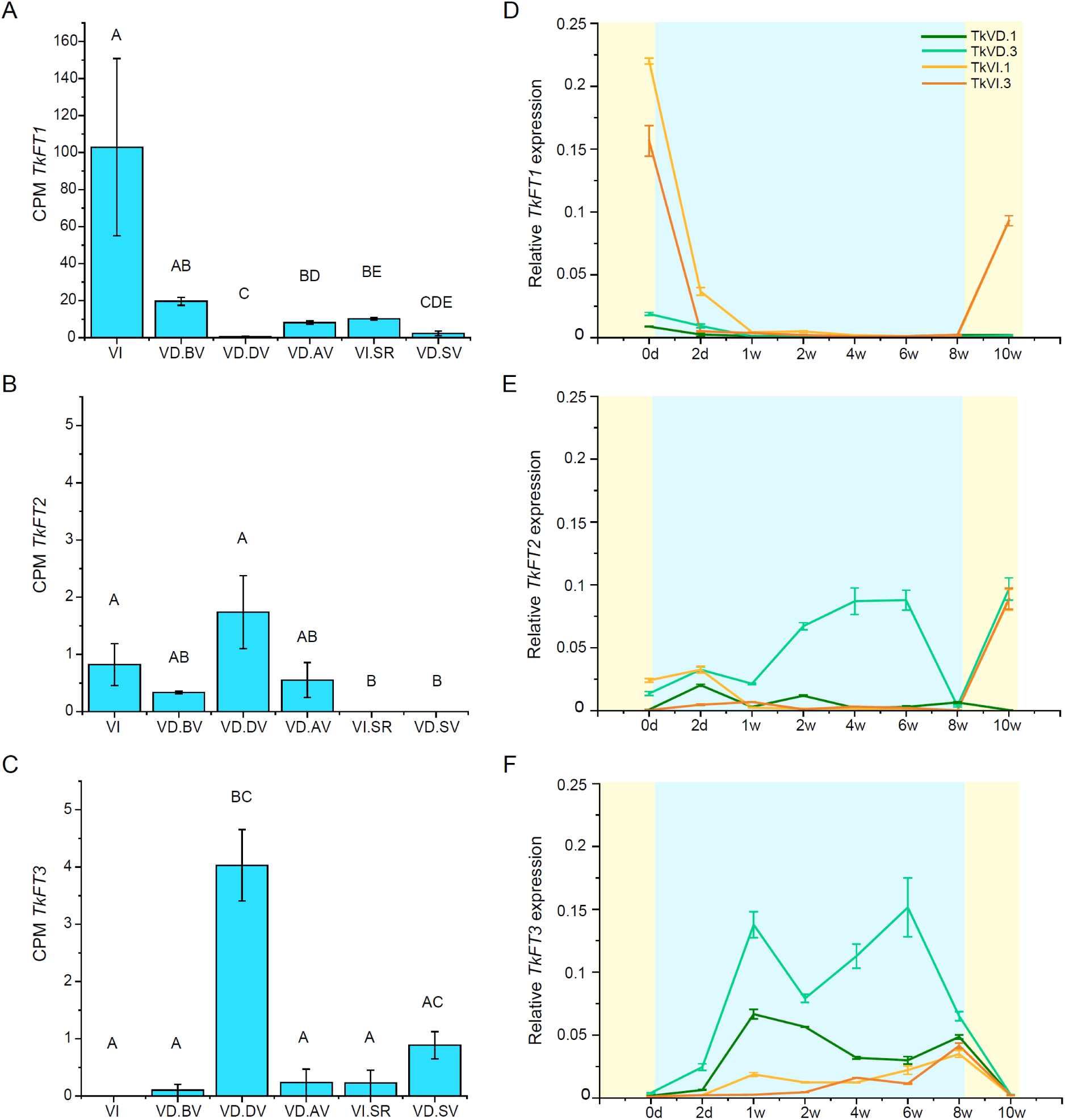
Expression levels of *TkFT1–3* in vernalization-independent (VI) and vernalization-dependent (VD) plants before (BV), during (DV) and after (AV) vernalization. **A)** *TkFT1*, **B)** *TkFT2* and **C)** *TkFT3* expression levels were analyzed by MACE in VI or VD plants using leaf tissue from VI, VD.BV, VD.DV, and VD.AV plants or shoot apical meristem enriched reproductive (VI.SR) and vegetative (VD.SV) tissue. Data are means ± SEM, *n* = 3 biological replicates (VI, VD.BV, VD.DV, VD.AV), *n* = 3 pools of five biological replicates (VI.SR, VD.SV). Different letters indicate statistically significant differences (FDR < 0.05; Benjamini-Hochberg). CPM = counts per million. **D-F)** Relative expression levels of *TkFT1–3* in VD (TkVD.1, TkVD.3) and VI *T. koksaghyz* (TkVI.1, TkVI.3) accessions before (0d) and during vernalization (2d, 1w, 2w, 4w, 6w, 8w) and after 8 weeks of vernalization and return to ambient temperature for 2 weeks (10w). Relative expression levels of D) *TkFT1*, E) *TkFT2* and F) *TkFT3* were determined by qRT-PCR using *TkRP* as a reference gene. Data are means ±SEM of technical triplicates, *n* = one biological replicate (TkVI.1, TkVI.3), *n* = pool of two biological replicates (TkVD.1, TkVD.3). Pale yellow background represents 22 °C, LD conditions and pale blue background represents 6 °C, SD conditions (d = days, w = weeks).

To mimic winter in the temperate climate zone, we typically apply SD conditions during vernalization. We therefore tested whether SD conditions were sufficient to inhibit *TkFT1* expression or induce *TkFT3* expression by analyzing diurnal expression profiles in pools of VI and VD plants under LD and SD conditions at ambient temperatures. *TkFT1* and *TkFT2* were expressed under LD conditions in VI and VD plants, whereas *TkFT3* expression was minimal (Fig. S3A-C). *TkFT1* expression was lower under SD than LD conditions in VI and VD plants except for the moderate expression observed in VD plants at ZT6 (Fig. S3D). *TkFT2* and *TkFT3* were minimally expressed under SD conditions (Fig. S3E, F). Accordingly, SD conditions alone are insufficient to abolish *TkFT1* expression or to induce *TkFT3* expression, as is the case during vernalization (Fig. 1A, C). Assuming that *FT* expression promotes the transition to flowering, the generally low levels of *TkFT1–3* expression under SD conditions at ambient temperatures are consistent with the finding that a 3-week SD treatment at ambient temperatures did not induce flowering in VD plants (Fig. S1B). However, 3 weeks of cold temperatures under LD or SD conditions induced flowering in VD plants (Fig. S1A, B), suggesting that vernalization is the most important factor promoting flowering in VD plants, whereas day length during the cold period is not critical for the flowering response.

To determine whether *TkFT1* and *TkFT3* expression in response to day length and temperature is associated with TFBSs in the corresponding promoters, we amplified and sequenced ∼1.6 kb of the promoter sequences from the genomic DNA of a VD accession (Tk203). We used the PlantPan 4.0 database (Chow et al. 2024) and Arabidopsis as the reference organism to analyze the promoter sequences *in silico*. After filtering low-similarity TFBSs (score < 0.8), 86 sites were detected, 71 of which were identified in the promoters of both *TkFT1* and *TkFT3* (Fig. S4, S5, Table S2). The transcription factors assumed to bind these sites in Arabidopsis mediate the cellular response to hormones, chromatin remodeling and/or regulation of reproductive process, among other functions. Interestingly, 22 transcription factors were found to be involved in the response to light, seven of which specifically regulate the photoperiodic control of flowering. Moreover, 12 transcription factors involved in cold responses, six involved in heat responses, and four involved in both were shown to recognize the corresponding TFBSs in Arabidopsis (Fig. S4, S5, Table S2). The promoters of both *TkFT1* and *TkFT3* thus contain potential TFBSs that could mediate photoperiodic or temperature responses.

### TkFT1–3 interact with TkFD1

FT proteins are known to form complexes with FD homologs mediated by 14-3-3 proteins (Taoka et al. 2011). Eudicot FDs share a conserved motif arrangement, namely the motifs A, LSL, bZIP and SAP (Tsuji et al. 2013). We identified one FD homolog with this motif arrangement in *T. koksaghyz*, and named it TkFD1 (Fig. S6A). MACE analysis indicated that *TkFD1* is predominately expressed in SAM-enriched tissue, which was verified by qRT-PCR using pooled material from corresponding samples (Fig. 3A, Fig. S6B). Its conserved domains and highly specific expression profile make TkFD1 an excellent candidate to be the cofactor of *T. koksaghyz* FT proteins. We studied the interaction between TkFT1–3 and TkFD1 by BIFC analysis based on the monomeric red fluorescent protein (mRFP1) (Jach et al. 2006). Confocal laser scanning microscopy revealed that TkFD1 interacted with all three TkFT proteins in the nucleus of *N. benthamiana* epidermal cells (Fig. 3B).

**Fig. 3.**
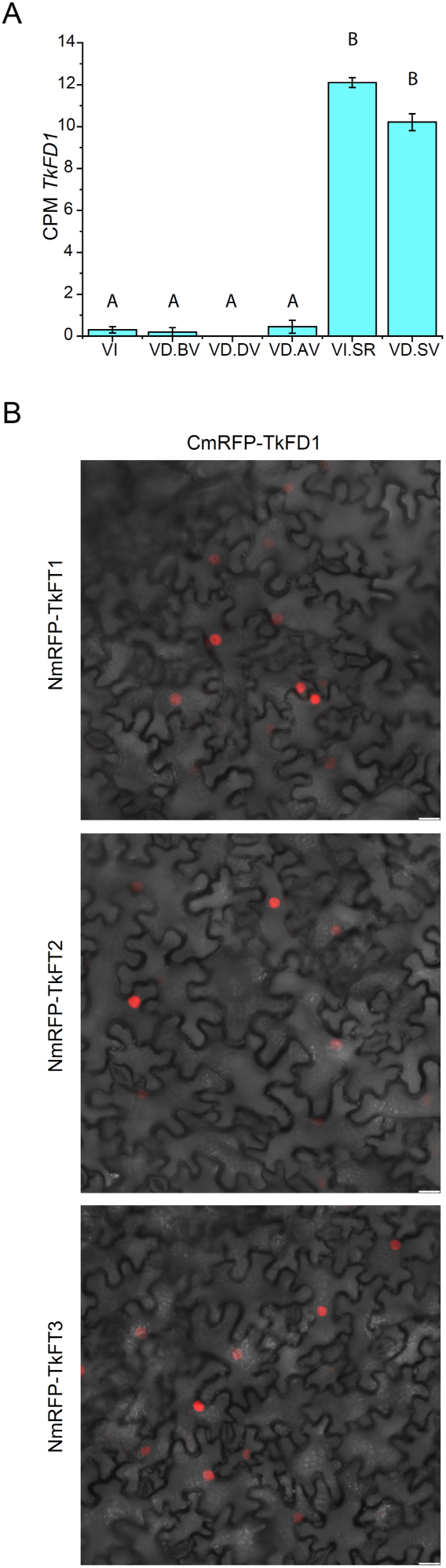
TkFD1 is expressed predominately in SAM-enriched tissue and interacts with TkFT1–3. **A)** *TkFD1* expression levels determined by MACE in VI or VD plants using leaf tissue (VI, VD.BV, VD.DV, VD.AV) or shoot apical meristem-enriched tissue (VI.SR, VD.SV). Data are means ± SEM, *n* = 3 biological replicates (VI, VD.BV, VD.DV, VD.AV), *n* = 3 pools of five biological replicates (VI.SR, VD.SV). Different letters indicate statistically significant differences (FDR < 0.05; Benjamini-Hochberg). VI = vernalization-independent, VD = vernalization-dependent, BV = before vernalization, DV = during vernalization, AV = after vernalization, SR = shoot apical meristem (reproductive), SV = shoot apical meristem (vegetative), CPM = counts per million. **B)** Bimolecular fluorescence complementation assay in *N. benthamiana* cells with TkFD1 and TkFT1–3. TkFD1 was C-terminally fused to the C-terminal mRFP1 fragment, and TkFT1–3 were C-terminally fused to the N-terminal mRFP1 fragment. Interaction was confirmed by reconstituted mRFP fluorescence visible in the nuclei. N-terminal or C-terminal mRFP1 fragments served as negative controls and were co-expressed with the corresponding TkFD1 or TkFT1– 3 fusion proteins. We did not observe nonspecific interactions in any combination. Scale bar = 20 μm.

### *TkFT1–3* overexpression promotes early flowering in transgenic plants

We investigated the ability of *TkFT1–3* to affect flowering time by cloning each coding sequence under the control of the PLsUbi promoter (including the 5′-UTR and the first intron), which confers strong and stable expression (Hirai et al. 2011; Kawazu et al. 2019). We transformed VD *T. koksaghyz* accession Tk203 to determine whether *TkFT1–3* overexpression was sufficient to overcome the need for vernalization (Fig. 4). Three of five regenerated T_0_ plants overexpressing *TkFT1* flowered in tissue culture, and one of these (line 4) also flowered after potting and repeatedly in the greenhouse (LD, 22 °C), correlating with particularly high *TkFT1* mRNA levels (Fig. 4B). One of three plants overexpressing *TkFT2* flowered 67 days (line 2) after potting, whereas the other two remained vegetative until the end of the experiment (200 days after potting) (Fig. 4C). Two of the three plants overexpressing *TkFT3* flowered 47 days (line 1) or 37 days (line 2) after potting (Fig. 4D). The expression levels of *TkFT2* and *TkFT3* in these plants reflected the flowering phenotype (Fig. 4C, D). As expected, the wild-type plants remained vegetative until the end of the experiment. All three TkFTs were thus able to bypass the vernalization requirement of *T. koksaghyz*. No T_1_ plants could be analyzed because *T. koksaghyz* is self-incompatible and a compatible crossing partner with a VD phenotype was unavailable. Therefore, we also overexpressed *TkFT1–3* in *T. officinale* and *T. brevicorniculatum*, both of which are triploid apomicts that produce clonal offspring. *T. officinale* undergoes VD flowering and *T. brevicorniculatum* undergoes VI flowering.

**Fig. 4.**
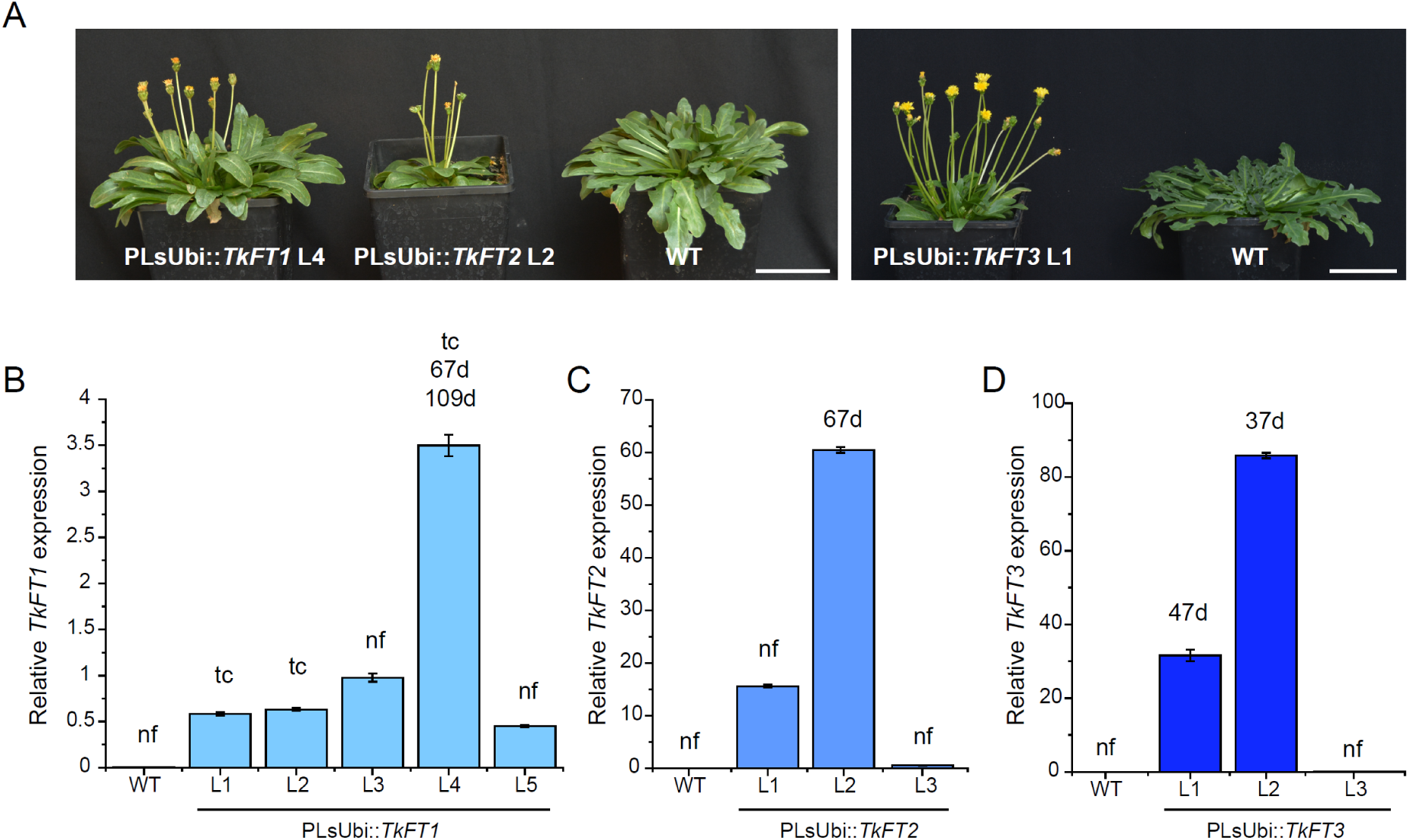
*TkFT1–3* overexpression bypasses the vernalization requirement in *T. koksaghyz*. **A)** Flowering phenotypes of *T. koksaghyz* Tk203 plants overexpressing *TkFT1* (PLsUbi::*TkFT1*), *TkFT2* (PLsUbi::*TkFT2*), or *TkFT3* (PLsUbi::*TkFT3*) and corresponding wild-type plants (WT) without vernalization. Scale bar = 6 cm. **B-D)** Relative *TkFT1–3* expression levels in the overexpression lines compared to the wild-type control, with indications of the flowering phenotype: nf = non-flowering at the end of the experiment (200 days after potting), tc = flowered in tissue culture, d = flowered in the greenhouse after the indicated number of days. Days until flowering were counted until the first floral bud had fully opened. Relative expression levels of B) *TkFT1*, C) *TkFT2* and D) *TkFT3* in the overexpression lines compared to wild-type controls were determined by qRT-PCR using *RP* as a reference gene. Data are means ±SEM of technical triplicates.

First, we transformed *T. officinale* to determine whether *TkFT1–3* overexpression is also sufficient to overcome the vernalization requirement in this species. The regenerated T_0_ plants were vernalized for 3 weeks to allow prompt seed production for the analysis of T_1_ progeny. We counted the days until flowering for two independent transgenic lines representing each construct along with wild-type plants as controls (Fig. S7). As expected, the wild-type plants had not flowered under the applied conditions (22 °C, LD) by the end of the experiment (200 days after sowing). Both lines overexpressing *TkFT1* were able to overcome the vernalization requirement. Line 1 flowered after 115 days and line 6 after 49 days, on average. One of the lines overexpressing *TkFT2* was able to overcome the vernalization requirement and flowered after 77 days on average, but the other remained vegetative until the end of the experiment. Both *TkFT3* overexpression lines were able to overcome the vernalization requirement and flowered after an average of 72 days for line1 or 94 days for line 2 (Fig. S7A). Higher *TkFT1*, *TkFT2* or *TkFT3* expression was strongly correlated with the flowering phenotype (Fig. S7, Table S3).

We transformed *T. brevicorniculatum* to determine whether *TkFT1–3* trigger early flowering in a VI species. We counted the days until flowering for two independent transgenic lines in the T_1_ generation for each construct along with vector control plants (Fig. 5A, B). *TkFT1* overexpression resulted in only a slight decrease in the number of days until flowering (∼3 days less than controls) in one of the two analyzed lines. In contrast, both lines overexpressing *TkFT2* flowered ∼11 days earlier than control plants. Plants overexpressing *TkFT3* showed the strongest phenotype, flowering 13 (line 4) or 15 (line 3) days earlier than controls (Fig. 5B). Higher *TkFT1*, *TkFT2* or *TkFT3* expression was significantly correlated with earlier flowering (Fig. 5B-E, Table S4). The combined overexpression data clearly demonstrate that all three TkFT proteins are potent floral inducers.

**Fig. 5.**
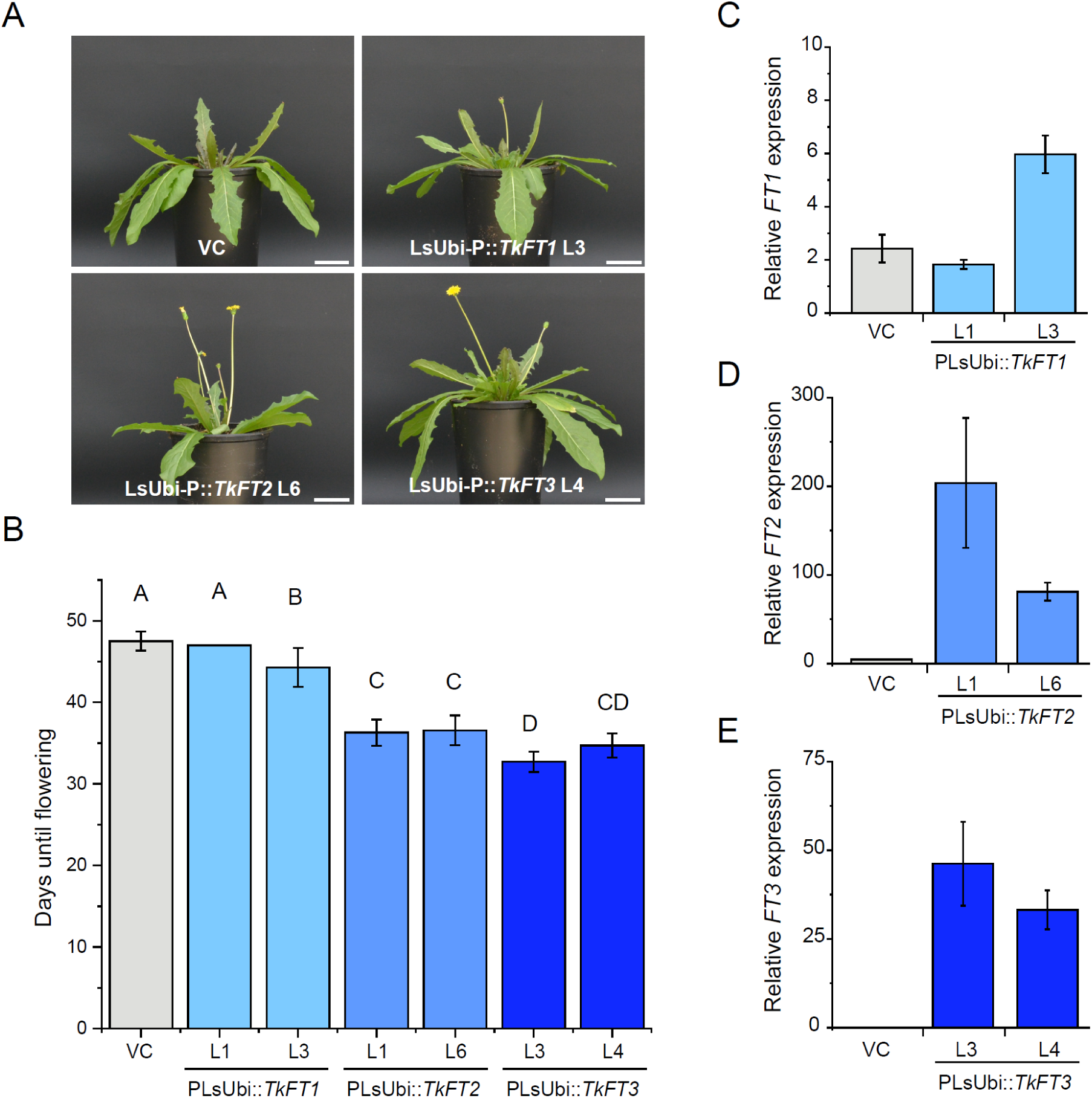
*TkFT1–3* overexpression promotes early flowering in *T. brevicorniculatum*. **A)** Flowering phenotype of *T. brevicorniculatum* overexpressing *TkFT1* (PLsUbi::*TkFT1*), *TkFT2* (PLsUbi::*TkFT2*) and *TkFT3* (PLsUbi::*TkFT3*) or vector control plants (VC). Pictures were taken 38 days after sowing. Scale bar = 6 cm. **B)** Days until flowering were counted until the first floral bud had fully opened. Data are means ± SD, *n* = 6 biological replicates (PLsUbi::*TkFT1*, PLsUbi::*TkFT2*, PLsUbi::*TkFT3*), *n* = 12 biological replicates (VC). Different letters indicate statistically significant differences as determined by ANOVA and Tukey’s *post hoc* test (p < 0.05). **C-E)** Relative expression levels of C) *TkFT1*, D) *TkFT2* and E) *TkFT3* in the overexpression lines compared to endogenous *TbFT* expression in VC determined by qRT-PCR using *RP* as a reference gene. Data are means ±SEM, *n* = 3 biological replicates.

### Two *FUL* homologs are downstream targets of TkFT1–3 in *Taraxacum* spp

In Arabidopsis, the MADS-box proteins AP1 and FUL are downstream targets of the florigen activation complex and they act redundantly to determine meristem identity, but have distinct roles in perianth identity and fruit development, respectively (McCarthy et al. 2015). In the Asteraceae species *Gerbera hybrida* and *Chrysanthemum morifolium*, FUL homologs have been shown to induce early flowering (Ruokolainen et al. 2010; Zhao et al. 2023). Based on RNA-Seq data, we identified two *T. koksaghyz* ORFs whose deduced amino acid sequences include a C-terminal FUL-like motif (Fig. S8A) – distinguishing them from AP1 proteins (Litt and Irish 2003) – and named them *TkFUL1* and *TkFUL2*. We analyzed their expression in VI and VD plants before, during and after vernalization, as well as in SAM-enriched tissue, by MACE and qRT-PCR (Fig. 6, Fig. S8B, C). *TkFUL1* was expressed at low levels in VI and VD plants before, during and after vernalization and also in reproductive and vegetative SAM-enriched tissue (Fig. 6A, Fig. S8B). *TkFUL2* was expressed in VI and VD plants before vernalization, was upregulated during vernalization, and its expression was maintained at rather high levels after vernalization. It was also expressed in both reproductive and vegetative SAM-enriched tissue (Fig. 6B, Fig. S8C). We next analyzed the expression of *TkFUL1* and *TkFUL2* (as well as their *T. officinale* and *T. brevicorniculatum* orthologs) in the *TkFT1–3* overexpression plants. The *FUL1* and *FUL2* genes were upregulated in all lines with elevated *TkFT* expression, confirming that *FUL1* and *FUL2* are downstream targets of TkFT1–3 (cf. Fig. 4B-D, Fig. 5C-E, Fig. S7B-D, Fig. 7).

**Fig. 6.**
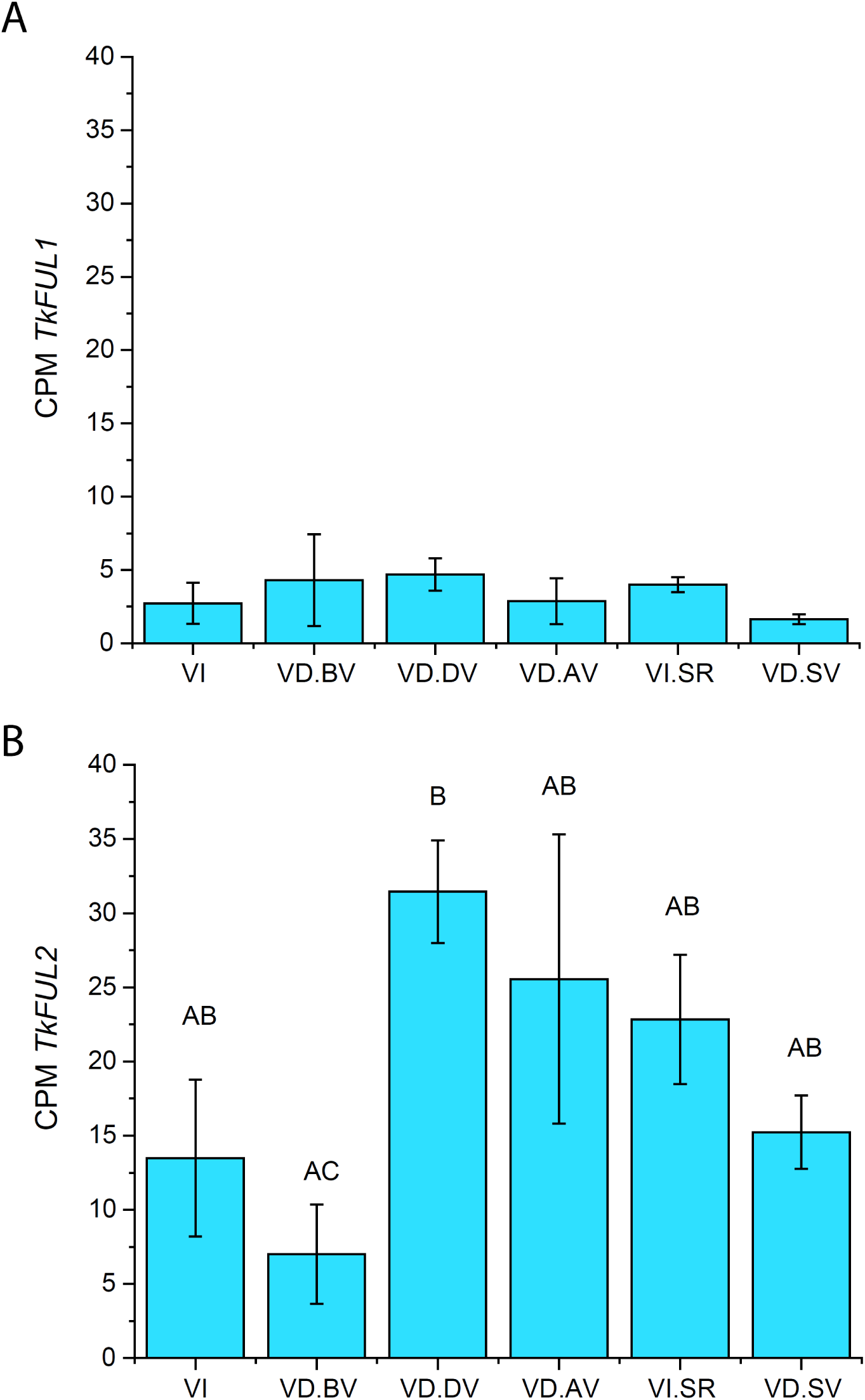
Expression levels of *TkFUL1* and *TkFUL2* in VI and VD *T. koksaghyz* before, during and after vernalization. **A)** *TkFUL1* and **B)** *TkFUL2* expression levels were analyzed by MACE in VI or VD plants using leaf tissue (VI, VD.BV, VD.DV, VD.AV) or shoot apical meristem-enriched tissue (VI.SR, VD. SV). Data are means ± SEM, *n* = 3 biological replicates (VI, VD.BV, VD.DV, VD.AV), *n* = 3 pools of five biological replicates (VI.SR, VD. SV). Different letters indicate statistically significant differences (FDR < 0.05; Benjamini-Hochberg). VI = vernalization-independent, VD = vernalization-dependent, BV = before vernalization, DV = during vernalization, AV = after vernalization, SR = shoot apical meristem (reproductive), SV = shoot apical meristem (vegetative), CPM = counts per million.

**Fig. 7.**
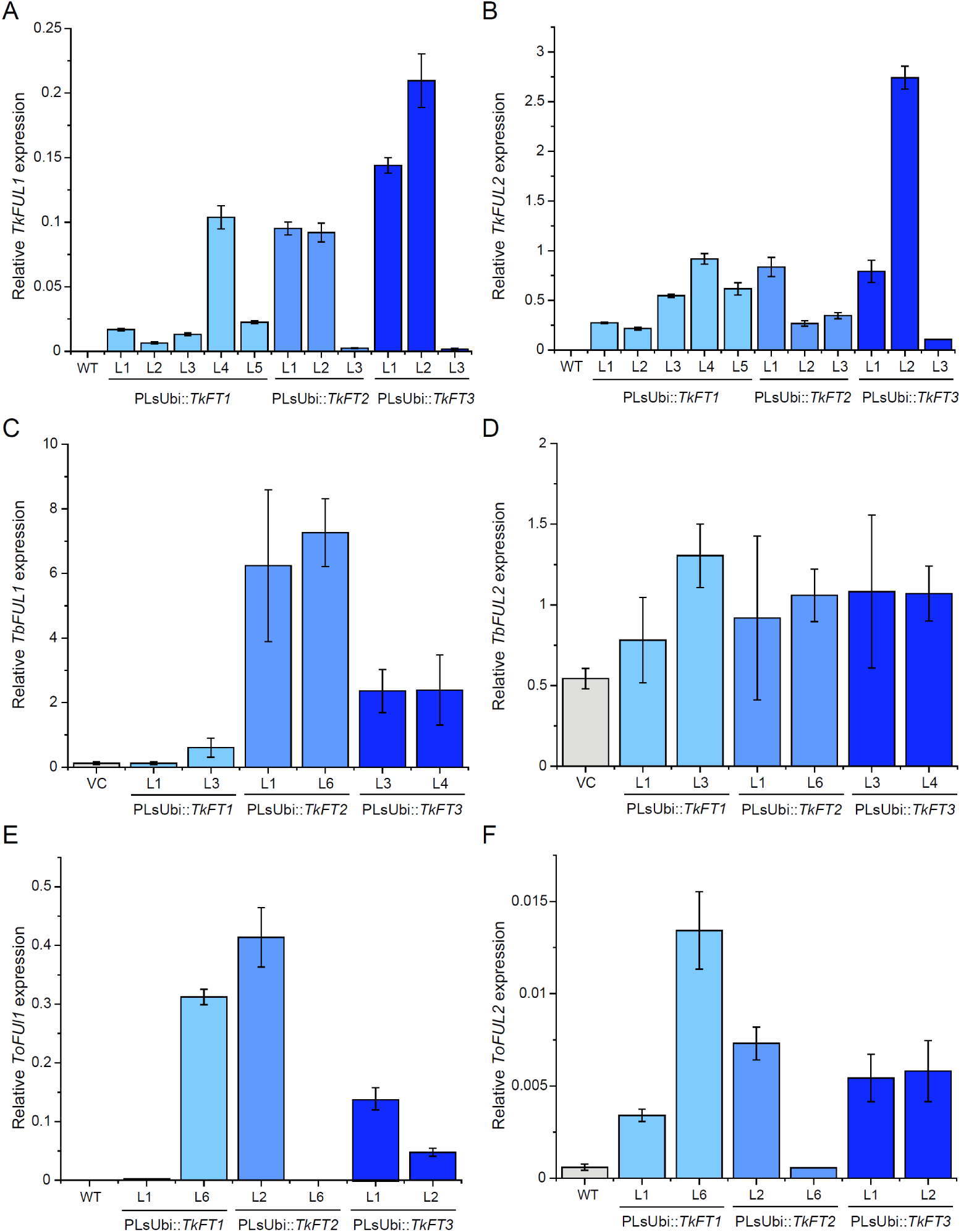
*FUL1* and *FUL2* are upregulated in *Taraxacum* spp. overexpressing *TkFT1–3*. **A-B)** Relative expression levels of A) *TkFUL1* and B) *TkFUL2* in *T. koksaghyz* Tk203 T_0_ plants overexpressing *TkFT1* (PLsUbi-P::*TkFT1*), *TkFT2* (PLsUbi::*TkFT2*) or *TkFT3* (PLsUbi::*TkFT3*) compared to endogenous *TkFUL1 and TkFUL2* expression in wild-type Tk203 (WT) determined by qRT-PCR using *RP* as a reference gene. Data are means ±SEM of technical triplicates. **C-D)** Relative expression levels of C) *TbFUL1* and D) *TbFUL2* in *T. brevicorniculatum* overexpressing *TkFT1* (PLsUbi-P::*TkFT1*), *TkFT2* (PLsUbi::*TkFT2*) or *TkFT3* (PLsUbi::*TkFT3*) compared to endogenous *TbFUL1 and TbFUL2* expression in vector controls (VC) determined by qRT-PCR using *RP* as a reference gene. Data are means ±SEM, *n* = 3 biological replicates. **E-F)** Relative expression levels of E) *ToFUL1* and F) *ToFUL2* in *T. officinale* overexpressing *TkFT1* (PLsUbi-P::*TkFT1*), *TkFT2* (PLsUbi::*TkFT2*) or *TkFT3* (PLsUbi::*TkFT3*) compared to endogenous *ToFUL1* and *ToFUL2* expression in wild-type (WT) plants determined by qRT-PCR using *RP* as a reference gene. Data are means ±SEM, *n* = 3 biological replicates

## Discussion

FT is a major network hub in flower development that integrates signals from several floral pathways and is conserved between species (Jin et al. 2021). *FT* expression is involved in the vernalization dependency of several species, including sugar beet and narrow-leafed lupin (*Lupinus angustifolius*). In biennial sugar beet, vernalization downregulates the floral repressor gene *BvFT1*, enabling the expression of the floral activator gene *BvFT2* during and after vernalization, ultimately leading to flowering (Pin et al. 2010; 2012). In narrow-leafed lupins, natural mutations (*Ku*, *Jul* and *Pal*) allow flowering without vernalization by deregulating the expression of the FTc1 protein (Nelson et al. 2017; Rychel-Bielska et al. 2020). FT homologs may also underpin the vernalization requirement of *Lupinus albus* (Rychel-Bielska et al. 2021).

In *T. koksaghyz,* vernalization and photoperiod orchestrate flowering in a complex manner. Without cold exposure, VI plants require LD conditions for floral induction, whereas exposure to low temperatures in the field during late fall/winter allows *T. koksaghyz* to flower under all photoperiod conditions (photoperiod 8–18 h). However, if the photoperiod is 12 h or longer, flowering occurs more readily (Borthwick et al. 1943; Krotkov 1945). This interdependence of vernalization and day length has been described as the “photothermal” induction of flowering (Owen et al. 1941).

The three *T. koksaghyz FT* genes (*TkFT1–3*) were shown to act as floral inducers: overexpression promoted flowering in *T. brevicorniculatum* and they were able to bypass the vernalization requirement in *T. koksaghyz* and *T. officinale*. Additionally, the TkFT1–3 proteins were shown to form florigen activation complexes with TkFD1, which is predominately expressed in SAM-enriched tissues. Finally, we identified *FUL* homologs that were upregulated by *TkFT1–3* overexpression in three *Taraxacum* species, indicating they are downstream targets of TkFT1–3.

At ambient temperatures, *TkFT1* is expressed at higher levels under LD than SD conditions, whereas *TkFT2* is expressed at significant levels exclusively under LD conditions (Fig. S3), aligning with the fact that VI flowering requires LD conditions (Krotkov 1945). The expression of the *TkFT* genes was also found to be sensitive to vernalization: *TkFT1* was significantly downregulated during vernalization, whereas *TkFT3* was expressed in VI and VD plants but only in response to cold exposure (Fig. 2, Fig. S2, S3), aligning with the fact that cold treatment promotes flowering in both cases (Krotkov 1945). *TkFT1* expression was upregulated again in VI plants after vernalization, supporting its role as major floral inducer in VI plants (Fig. 2D). In contrast, *TkFT1* expression in VD plants did not increase after vernalization relative to pre-vernalization levels, suggesting TkFT1 has no major role in floral induction in VD plants after vernalization (Fig. 2A, D).

Vernalization in its true sense is a preparatory process that creates flowering competence, which becomes noticeable only following the initiation of floral primordia and eventually flower development (Chouard 1960). The strong expression of the floral inducer *TkFT3* during vernalization suggests that it has a major role during this preparatory process by supporting the capacity for subsequent flowering. We found not only that *TkFT3* is induced during vernalization but also its downstream target *TkFUL2*, and remarkably, its elevated expression level persists afterward (Fig. 6B; Fig. S8C).

Further studies are needed to identify the genetic factors controlling the specific upregulation of *TkFT3* during vernalization, and with emphasis on the general role of *FT* genes, the expression of *TkFT1–3* in VI and VD plants under varying environmental conditions. Recently, we identified promising candidate genes involved in the control of flowering time in *T. koksaghyz* (Roelfs et al. 2024). Interestingly, Arabidopsis homologs of some of those candidates, including homologs of EARLY FLOWERING (ELF) and CHROMATIN REMODELING (CHR) (Table S2, Fig. S4, S5), recognize putative TFBSs in the *TkFT1* and *TkFT3* promoters (Roelfs et al. 2024). To better understand this complex regulatory network, upstream factors and downstream targets of the hub proteins TkFT1–3 should be characterized in more detail.

In summary, our findings indicate that *TkFT1* expression in VI plants is sufficient to induce downstream target genes such as *TkFUL1* and *TkFUL2* under ambient temperatures and LD conditions. *TkFT3* expression during vernalization in VI plants may have an additive effect, leading to earlier flowering after cold treatment, an observation that has been described previously (Krotkov 1945). In VD plants, TkFT1 probably plays no major role in flowering after vernalization. Instead, high levels of *TkFT3* expression during vernalization are necessary to maintain or even upregulate the expression of downstream target genes such as *TkFUL1* and *TkFUL2* during and after vernalization, ultimately enabling flowering after vernalization. *TkFT2* probably has additive effects on both VI and VD flowering because it is moderately expressed in VI and VD plants under all tested conditions except generally non-inductive SD conditions at ambient temperatures (Fig. 2B, E; Fig. S3B, E). Our results clearly demonstrate that TkFT1 and TkFT3 make distinct contributions to VD and VI floral transition. Understanding how they are regulated and which target genes they control will reveal the molecular basis of vernalization dependency in *T. koksaghyz* in unprecedented detail. The same genes could then be targeted in domestication and breeding programs, allowing *T. koksaghyz* to be developed as an alternative crop for the production of natural rubber and other valuable metabolites.

## Acknowledgements

The authors would like to thank Dr. Fred Eickmeyer (ESKUSA GmbH, Germany) for the provision of *T. koksaghyz* plants. We also thank Dr. Hiroyuki Fukuoka (Takii Plant Breeding and Experiment Station, Japan) for the PLsUbi sequence. We thank Martina Koch and Sascha Ahrens (both University of Münster), Andreas Wagner and Dr. Jost Muth (both Fraunhofer IME, Aachen) for technical and horticultural assistance.

## Statements and Declarations

### Author contributions

Conceptualization, A.K., G.A.N., D.P.; investigation, A.K., K.U.R., M.W., B.L., M.K.; validation, A.K., K.U.R; M.W.; data analysis, A.K., K.U.R., B.L.; visualization, A.K., K.U.R.; writing – original draft, A.K., K.U.R.; writing – review & editing, A.K., R.M.T., G.A.N., D.P.; funding acquisition, D.P.; project administration, A.K., G.A.N., D.P., supervision: D.P. All authors read and approved the final manuscript.

### Data availability

The authors declare that all data presented in this report are included within the paper or in the supplementary tables, figures, or text, or in publicly available databases.

