## Supplementary text for "Temperature-regulated *FLOWERING LOCUS T* homologs make distinct contributions to the floral transition in vernalization-dependent and vernalization-independent *Taraxacum koksaghyz* plants": Supplementary text.docx

**Title:**

**Author information:**

Andrea Känel^1‡^, Kai-Uwe Roelfs^1^, Michael Wissing^2^, Benjamin Lenzen^1^, Malin Klein^1^, Richard M. Twyman^3^, Gundula A. Noll^1,2^, Dirk Prüfer^1,2^

^1^ Institute of Plant Biology and Biotechnology, University of Muenster, Muenster, Germany

^2^ Fraunhofer Institute for Molecular Biology and Applied Ecology IME, Muenster, Germany

^3^ TRM Ltd, Scarborough, UK

^‡^ Corresponding author:

Andrea Känel

**Supplementary text**

Protein IDs of *T. koksaghyz* PEBP homologs according to Lin et al. (2022)

TkMFT1: GWHPBCHF029259; TkMFT2: GWHPBCHF029225; TkBFT1: GWHPBCHF018972;

TkBFT2: GWHPBCHF037464; TkTFL1: GWHPBCHF054490; TkFT1: GWHPBCHF014458; TkFT2: GWHPBCHF032201

Deduced amino acid sequence of *TkFT3* isolated from *T. koksaghyz* Tk203

>TkFT3

MSRERDPLVIGRVIGDVLDGFTKSINLSVTYNDRVVNNGCELKPSQVVNQPRADIGGDDLRAFHTLVMVDPDAPSPSNPNLREYLHWLVTDIPATTGARFGQEVVCYESPRPSMGIHRLAFVLFRQMGRQTVYAPGWRQNFNTKDFAELYNLGSPVAAVYFNCQRESGLGGRRR

Protein IDs of FD proteins are available at NCBI (www.ncbi.nlm.nih.gov)

NtFD3: AVG70950.1; AtFD: NP_195315.3; SlSGPB: NP_001234345.1

Deduced amino acid sequence of *TkFD1* isolated from *T. koksaghyz* Tk203

>TkFD1

MLSDGGGCGGDDVNKNLNTSHHNLGRVSTRSTTLSSSSSSNTSSNHSPFSVSLTVNLIKS

TPTATRTMEEVWKDINLCSTTNHPAAGTKGYRGFILQDFLAKPFSNNETPTSLSSPGYGS

PSPPPPAPPTSQPSMLINLNSGPDQLNFLADDPTRNVSPLDGPFDHVLGSSNSSSGLLGQ

SIGGMRMLPATDRVGGGERRHKRMIKNRESAARSRARKQAYMNELENEVERLKEENAKLK

RQQQELQVAPAAQMTKKGGLQRTSTAPF

Protein IDs of AP1-like and FUL-like proteins are available at NCBI (www.ncbi.nlm.nih.gov)

FUL-like sequences:

CmFL1: WHS04460.1; GSQUA5: CAX65663.1; GSQUA2: CAX65661.1; AtFUL: sp|Q38876.1

AP1-like sequences:

AtAP1: sp|P35631.2; AtCAL: sp|Q39081.3; AmSQUA: sp|Q38742; GSQUA3: CAX65662.1; CmAP1: AOV18945.1

Deduced amino acid sequences of *TkFUL1* and *TkFUL2* isolated from *T. koksaghyz* Tk203

>TkFUL1

MGRGRVQLKRIENKISRQVTFSKRRTGLLKKAHEISVLCDAHVALIVFSSRGKLFEYSNNSSMEAILERY

ERCAYAEKLLNGPETETPGSWTLESSQLMAKVEVLEKTIRHYVGEGLESLNLRELQNVEQHLDTALKRIR

TKKNQVMNESISQLHKKEKTLQEQRNTLYKKLKETEGNTNTTEPETLPINRDSFMESNLREEEYAGGHHI

TAAQLPPWMLQHVHQ

>TkFUL2

MGRGRVTLKRIENKINRQVTFSKRRSGLLKKAHEISVLCDADVALIVFSTKGKLCEYASDASMERILERH

ERHSYTERQLNGTDPQSQENWSLEHAKLKARIELFQKNQRHLMGEDLDSLSLKELQNYEQQLDTALRRLR

LRKNQLMLESISDLQKKDKALQDQNNLLLKEMKEKEKEVPQLPPMIIEQQTHDNIGTLNLGEMYQAGGDG

EIEDTRRQVMPHWILQYMNQ
