## Supplementary Figures S1-S8 for "Temperature-regulated *FLOWERING LOCUS T* homologs make distinct contributions to the floral transition in vernalization-dependent and vernalization-independent *Taraxacum koksaghyz* plants": Supplementary_Figures_S1-S8.DOCX

**Title:**

**Author information:**

Andrea Känel^1‡^, Kai-Uwe Roelfs^1^, Michael Wissing^2^, Benjamin Lenzen^1^, Malin Klein^1^, Richard M. Twyman^3^, Gundula A. Noll^1,2^, Dirk Prüfer^1,2^

^1^ Institute of Plant Biology and Biotechnology, University of Muenster, Muenster, Germany

^2^ Fraunhofer Institute for Molecular Biology and Applied Ecology IME, Muenster, Germany

^3^ TRM Ltd, Scarborough, UK

^‡^Corresponding author:

Andrea Känel

**Supplementary figures**

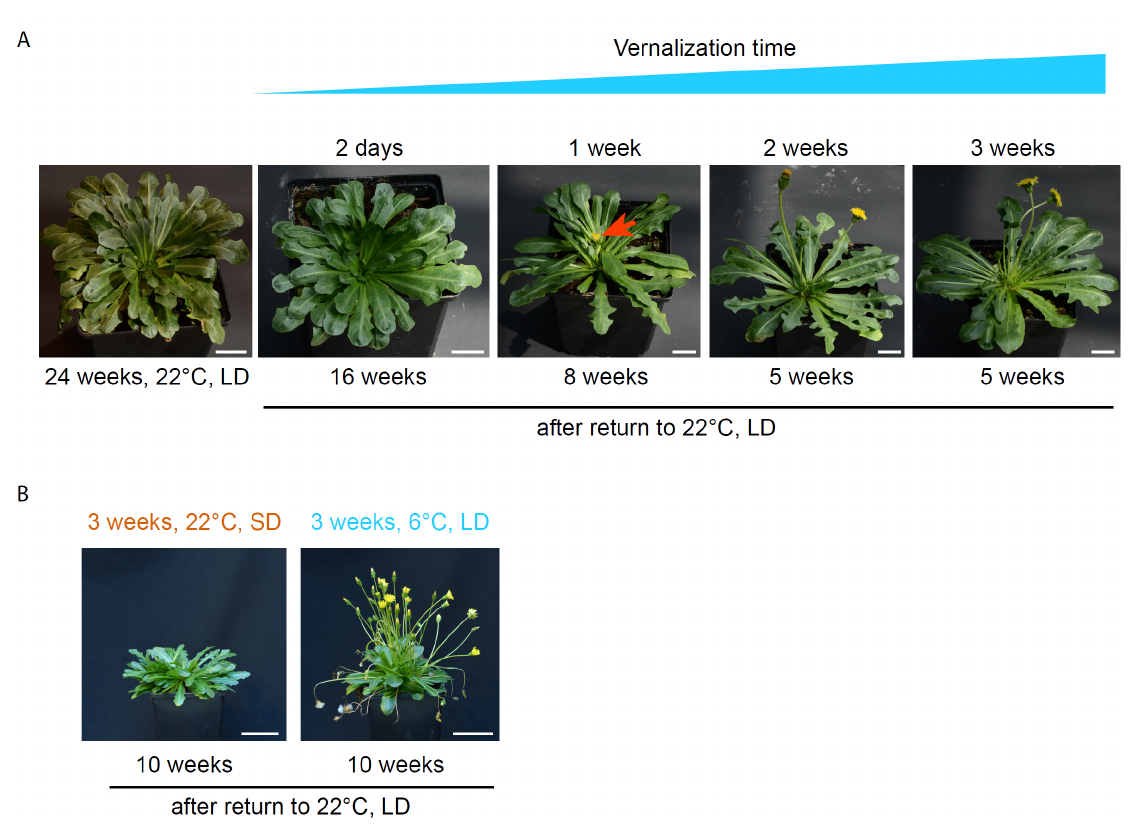

**Fig. S1** Flowering phenotype of vernalization-dependent (VD) *T. koksaghyz* under different cultivation conditions. **A)** VD *T. koksaghyz* accession Tk203 was cultivated for 8 weeks at 22 °C under long-day (LD) conditions in the greenhouse. Plants were then vernalized at 6 °C and short-day (SD) conditions for 2 days, 1 week, 2 weeks or 3 weeks. Afterwards, the plants were transferred back to LD conditions at 22 °C and the flowering phenotype was determined. Days until flowering were counted until the first floral bud had fully opened. One week of vernalization was sufficient to induce flowering after 8 weeks in five of six biological replicates. Extension of the vernalization time to ≥ 2 weeks decreased the time until flowering to 5 weeks (*n* = 6 biological replicates). Scale bar = 2 cm **B)** *T. koksaghyz* accession Tk203 was cultivated for 8 weeks under LD conditions at 22 °C in the greenhouse. Plants were then cultivated under SD conditions at 22 °C or LD conditions at 6 °C for 3 weeks and then transferred back to 22 °C under LD conditions (*n* = 3 biological replicates). Plants exposed to 6 °C and LD conditions induced flowering, whereas those exposed to 22 °C and SD conditions failed to induce flowering. Scale bar = 6 cm.

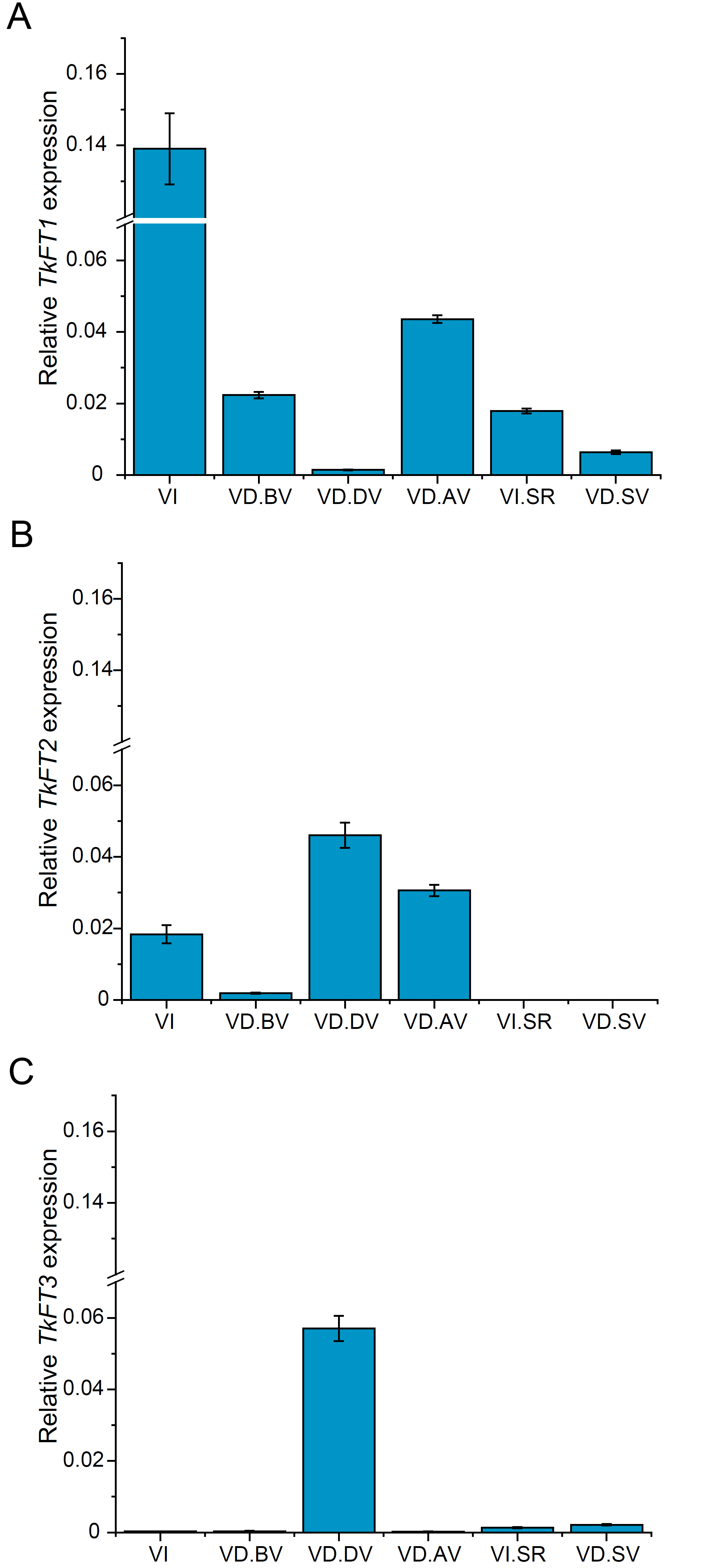

**Fig. S2** Relative expression levels of *TkFT1-3* in pools of VI and VD plants before, during and after vernalization. Relative expression levels of **A)** *TkFT1*, **B)** *TkFT2* and **C)** *TkFT3* were determined by qRT-PCR using *TkRP* as a reference gene. Data are means ±SEM, *n* = pool of 99 biological replicates (VI), *n* = pool of 104 biological replicates (VD.BV), *n* = pool of 39 biological replicates (VD.DV, VD.AV), *n* = pool of 33 biological replicates (VI.SR), *n* = pool of 27 biological replicates (VD.SV). VI = vernalization-independent, VD = vernalization-dependent, BV = before vernalization, DV = during vernalization, AV = after vernalization, SR = shoot apical meristem (reproductive), SV = shoot apical meristem (vegetative).

**Fig. S3** Diurnal expression profiles of *TkFT1–3* in VD and VI *T. koksaghyz* plants. **A-C)** *TkFT1–3* expression levels in VD (TkVD.1, TkVD.3 and Tk203) and VI (TkVI.1, TkVI.3) *T. koksaghyz* accessions grown under long-day (LD) or **D-F)** short-day (SD) conditions at ambient temperature. Relative expression values for A, D) *TkFT1*; B, E) *TkFT2*; and C, F) *TkFT3* determined by qRT-PCR using *TkRP* as a reference gene. Data are means ±SEM of technical triplicates, *n* = pool of three biological replicates. Leaves were harvested at the indicated Zeitgeber time (ZT 0 = beginning of light phase). White background indicates day time and gray background indicates night time. VI = vernalization-independent, VD = vernalization-dependent.

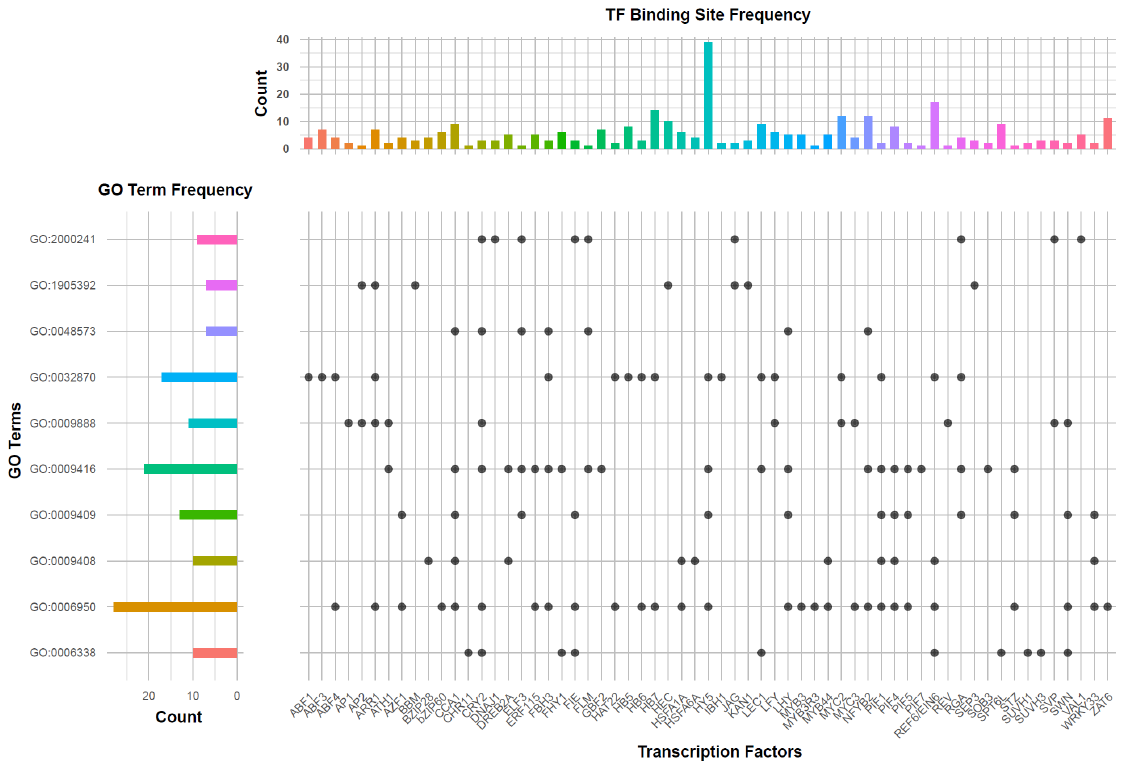

**Fig. S4** Visualization of the frequency of identified transcription factor binding sites (TFBSs) in the *TkFT1* promoter sequence that are known to be bound by the indicated transcriptions factors (TFs) in Arabidopsis, and selected GO annotations thereof. The upper bar plot indicates the frequency of identified TFBSs for each TF. The left bar plot indicates the frequency of TFs for the corresponding GO terms. The scatter plot maps the listed TFs on the lower axis to GO terms: response to heat (GO:0009408), response to cold (GO:0009409), response to light stimulus (GO:0009416), photoperiodism, flowering (GO:0048573), cellular response to hormone stimulus (GO:0032870), chromatin remodeling (GO:0006338), plant organ morphogenesis (GO:1905392), tissue development (GO:0009888), regulation of reproductive process (GO:2000241), and response to stress (GO:0006950).

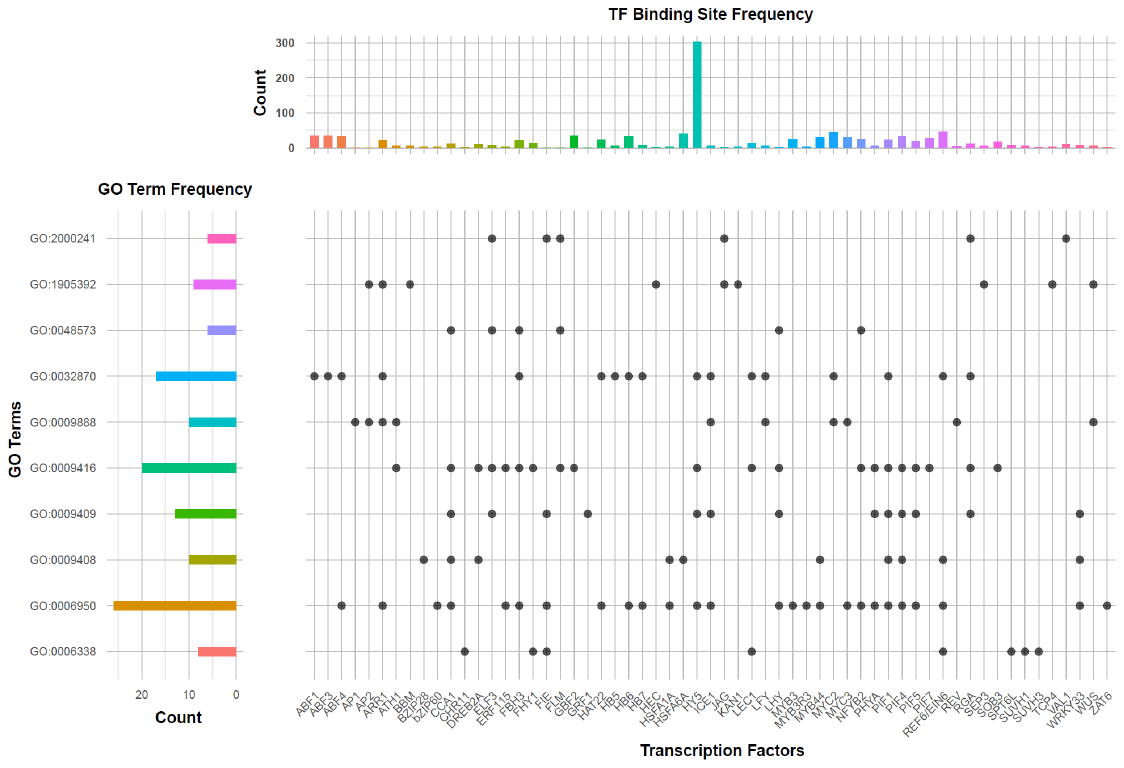

**Fig. S5** Visualization of the frequency of identified transcription factor binding sites (TFBSs) in the *TkFT3* promoter sequence that are known to be bound by the indicated transcriptions factors (TFs) in Arabidopsis, and selected GO annotations thereof. The upper bar plot indicates the frequency of identified TFBSs for each TF. The left bar plot indicates the frequency of TFs for the corresponding GO terms. The scatter plot maps the listed TFs on the lower axis to GO terms: response to heat (GO:0009408), response to cold (GO:0009409), response to light stimulus (GO:0009416), photoperiodism, flowering (GO:0048573), cellular response to hormone stimulus (GO:0032870), chromatin remodeling (GO:0006338), plant organ morphogenesis (GO:1905392), tissue development (GO:0009888), regulation of reproductive process (GO:2000241), and response to stress (GO:0006950).

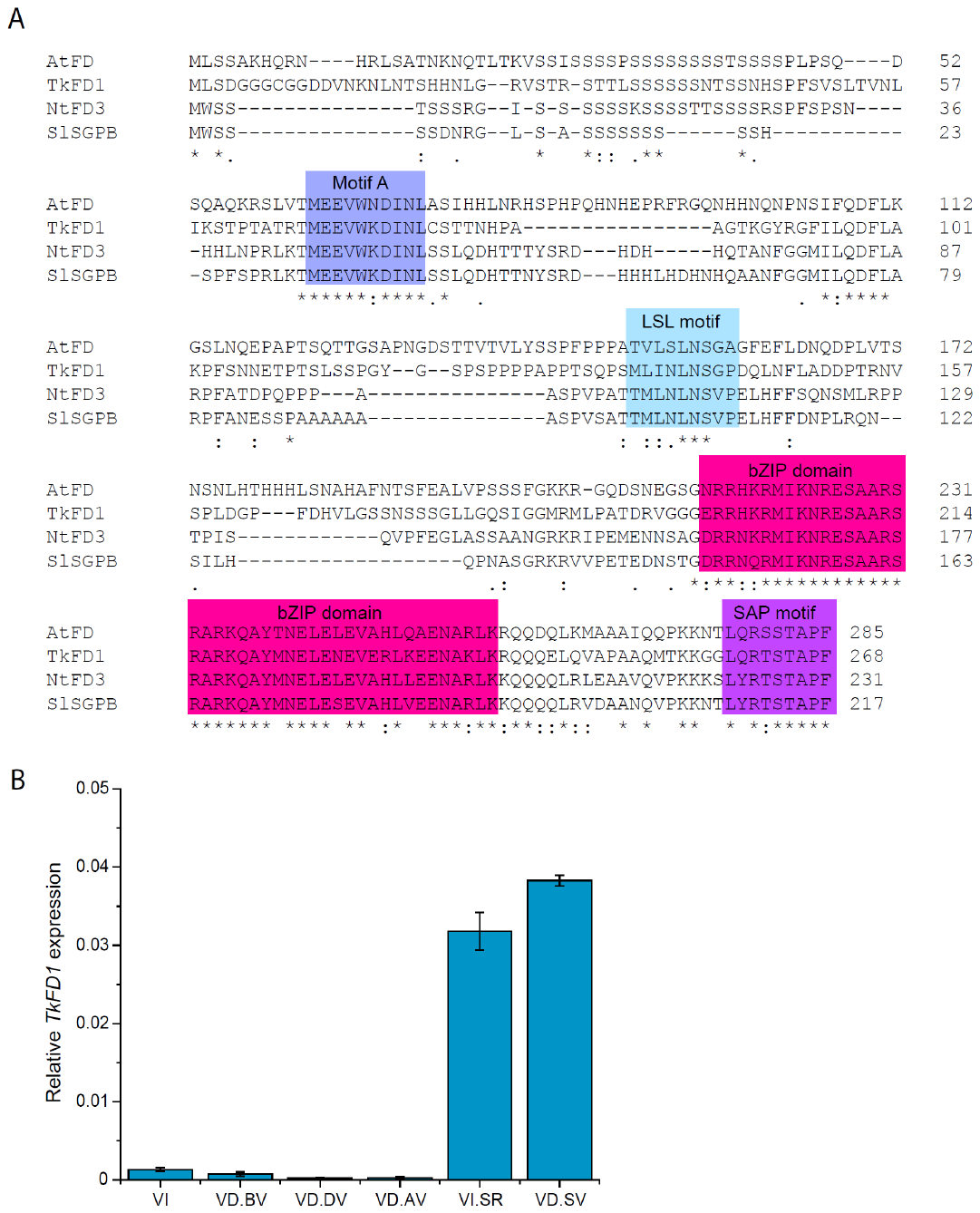

**Fig. S6** TkFD1 contains motifs that are shared by eudicot FD proteins and is mostly expressed in SAM-enriched tissue. **A)** The amino acid sequence alignment of TkFD1 with functionally validated FD proteins from Arabidopsis (AtFD), tobacco (NtFD3; Beinecke et al., 2018), and tomato (SPGB; Pnueli et al., 2001) shows that TkFD1 contains all four domains shared by eudicot FD proteins (motifs A, LSL, bZIP, and SAP). The multiple sequence alignment was created using the Clustal Omega program (Madeira et al. 2024). **B)** Relative expression levels of *TkFD1* were determined by qRT-PCR using *TkRP* as a reference gene. Data are means ±SEM, *n* = pool of 99 biological replicates (VI), *n* = pool of 104 biological replicates (VD.BV), *n* = pool of 39 biological replicates (VD.DV, VD.AV), *n* = pool of 33 biological replicates (VI.SR), *n* = pool of 27 biological replicates (VD.SV). VI = vernalization-independent, VD = vernalization-dependent, BV = before vernalization, DV = during vernalization, AV = after vernalization, SR = shoot apical meristem (reproductive), SV = shoot apical meristem (vegetative).

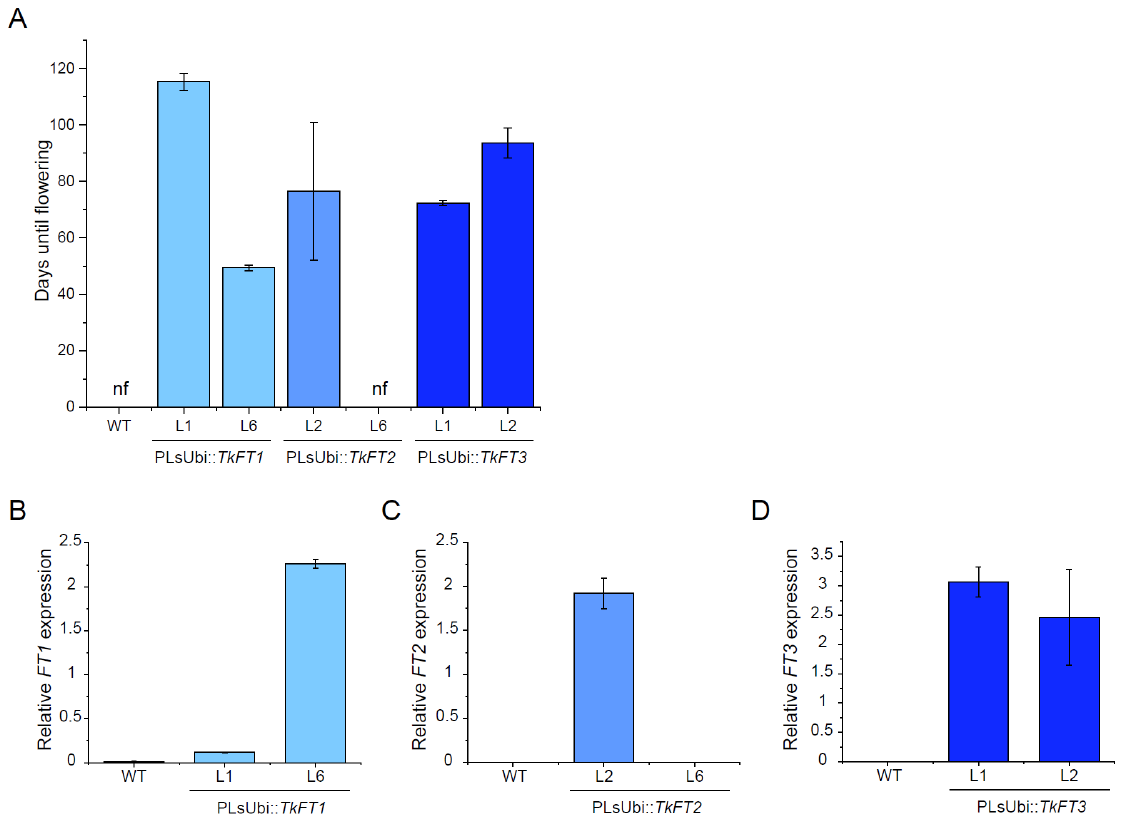

**Fig. S7** *TkFT1–3* overexpression overcomes the vernalization requirement in *T. officinale.* **A)** Days until flowering in *T. officinale* plants overexpressing *TkFT1* (PLsUbi::*TkFT1*), *TkFT2* (PLsUbi::*TkFT2*), or *TkFT3* (PLsUbi::*TkFT3*) and wild-type (WT) controls were counted until the first floral bud had fully opened. Data are means ± SD, *n* = 6 biological replicates: nf = non-flowering at the end of the experiment (200 days after sowing). **B-D)** Relative expression levels of B) *TkFT1*, C) *TkFT2* and D) *TkFT3* in the transgenic lines compared to endogenous *ToFT* expression in WT controls determined by qRT-PCR using *RP* as a reference gene. Data are means ±SEM, *n* = 3 biological replicates.

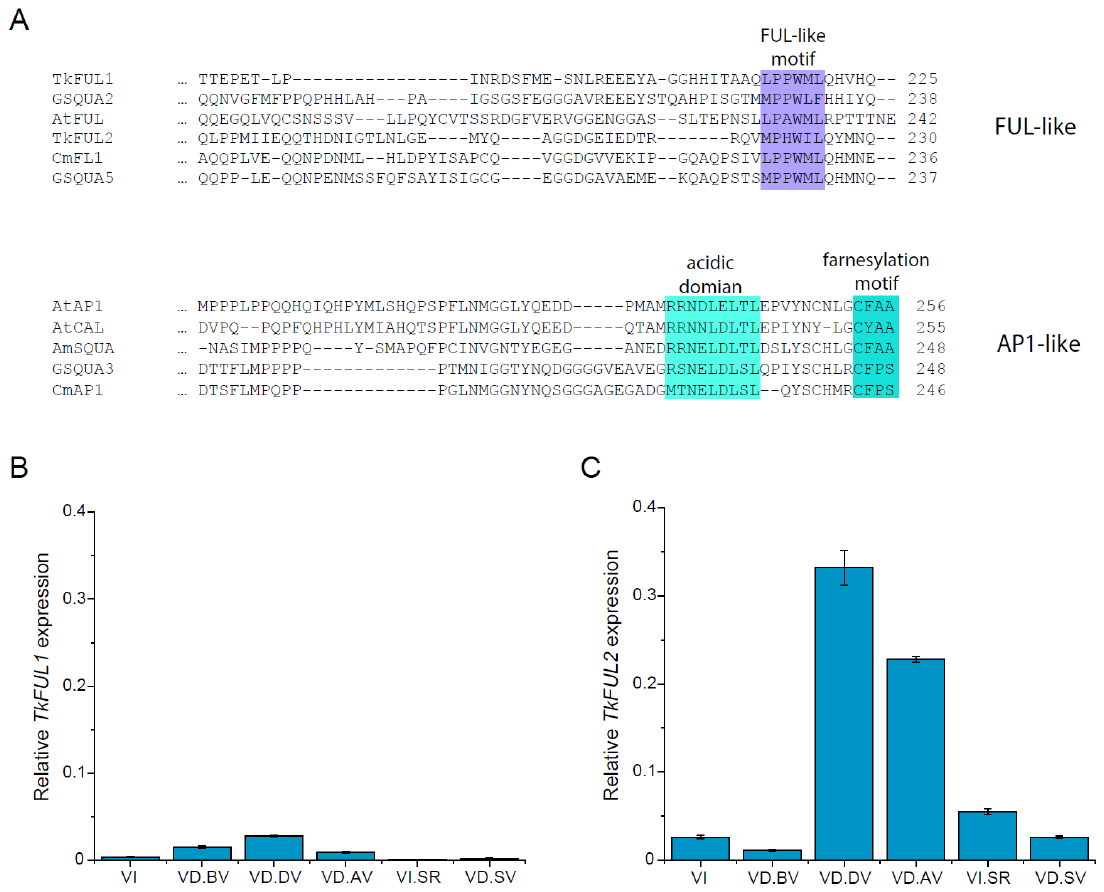

**Fig. S8** TkFUL1 and TkFUL2 belong to the FRUITFULL-like clade of the AP1/FUL lineage of MADS box proteins. **A)** TkFUL1 and TkFUL2 contain a C-terminal FUL-like motif. Proteins belonging to the AP1-like clade possess an acidic transcriptional activator domain (Cho et al., 1999) and a farnesylation motif. The multiple sequence alignment was created using the Clustal Omega program (Madeira et al. 2024). **B-C)** Relative expression levels of B) *TkFUL1* and C) *TkFUL2* were determined by qRT-PCR using *TkRP* as a reference gene. Data are means ±SEM, *n* = pool of 99 biological replicates (VI), *n* = pool of 104 biological replicates (VD.BV), *n* = pool of 39 biological replicates (VD.DV, VD.AV), *n* = pool of 33 biological replicates (VI.SR), *n* = pool of 27 biological replicates (VD.SV). VI = vernalization-independent, VD = vernalization-dependent, BV = before vernalization, DV = during vernalization, AV = after vernalization, SR = shoot apical meristem (reproductive), SV = shoot apical meristem (vegetative).
